# Starvation resistance is associated with developmentally specified changes in sleep, feeding, and metabolic rate

**DOI:** 10.1101/397455

**Authors:** Elizabeth B. Brown, Melissa E. Slocumb, Milan Szuperak, Arianna Kerbs, Allen G. Gibbs, Matthew S. Kayser, Alex C. Keene

## Abstract

Food shortage represents a primary challenge to survival, and animals have adapted diverse developmental, physiological, and behavioral strategies to survive when food becomes unavailable. Starvation resistance is strongly influenced by ecological and evolutionary history, yet the genetic basis for the evolution of starvation resistance remains poorly understood. The fruit fly, *Drosophila melanogaster*, provides a powerful model for leveraging experimental evolution to investigate traits associated with starvation resistance. While control populations only live a few days without food, selection for starvation resistances results in populations that can survive weeks. We have previously shown that selection for starvation resistance results in increased sleep and reduced feeding in adult flies. Here, we investigate the ontogeny of starvation resistance-associated behavioral and metabolic phenotypes in these experimentally selected flies. We find that selection for starvation resistance results in delayed development and a reduction in metabolic rate in larvae that persists into adulthood, suggesting that these traits may allow for the accumulation of energy stores and an increase in body size within these selected populations. In addition, we find that sleep is largely unaffected by starvation- selection and that feeding increases during the late larval stages, suggesting that experimental evolution for starvation resistance produces developmentally specified changes in behavioral regulation. Together, these findings reveal a critical role for development in the evolution of starvation resistance and indicate that selection can selectively influence behavior during defined developmental timepoints.

**SUMMARY STATEMENT:** *Drosophila melanogaster* selected for starvation resistance take longer to develop and exhibit development-specific changes in traits associated with the accumulation and conservation of energy stores.

## INTRODUCTION

Food acquisition represents a major challenge to many animal species, and the ability to locate food, or survive in the absence of food, strongly associates with reproductive fitness (Chippindale et al., 1996; Wayne et al., 2006). Starvation resistance varies dramatically throughout the animal kingdom, and even between closely related species, yet surprisingly little is known about the biological basis for evolved differences in this behavior (Gibbs and Reynolds, 2012; Matzkin et al., 2009; Rion and Kawecki, 2007). Animals have developed diverse mechanisms for responding to acute shortages in nutrient availability, including the induction of foraging behavior, alterations in sleep and locomotor activity, and changes in metabolic rate (Schmidt, 2014; Stahl et al., 2017; Sternson et al., 2013; Yurgel et al., 2014). While starvation resistance is likely influenced by developmental processes that contribute to an organism’s size, metabolic phenotypes, and brain function, it is not known whether selection occurs at developmentally specified stages or is maintained throughout development. Defining the effects of selection for starvation resistance on behavior and metabolism across development is therefore critical for understanding the developmental specificity of evolved changes in these processes.

The fruit fly, *Drosophila melanogaster* provides a powerful model for investigating the mechanistic basis of starvation resistance (Rion and Kawecki, 2007). Outbred populations of fruit flies display highly variable starvation resistance, as well as traits that are associated with starvation resistance including developmental timing, sleep, and feeding behaviors (Folguera et al., 2008; Garlapow et al., 2016; Harbison et al., 2017; Masek et al., 2014; Svetec et al., 2015; Yadav and Sharma, 2014), but little is known about how these individual traits contribute to the evolution of starvation resistance. We have implemented experimental evolution by starving outbred adult *Drosophila* until only 15% of the initial population remain alive, then passaging the survivors onto the next generation (Hardy et al., 2018). These populations have been independently selected over 100 generations resulting in flies that survive up to two weeks in the absence of food, while non-selected flies survive for only 3-4 days. These starvation-selected populations provide an opportunity to examine how behavioral and physiological traits are altered by selection for starvation resistance, and whether selection in adults also influences their development.

Altered life history and behavioral changes in adults are associated with evolutionarily acquired resistance to nutrient stress (Bubliy and Loeschcke, 2005; Gefen, 2006; Kolss et al., 2009), but the specific contributions of the many behavioral and physiological changes to starvation resistance has been difficult to test experimentally. We have previously identified increased sleep and reduced feeding in adult *Drosophila* selected for resistance to starvation stress (Masek et al., 2014; Slocumb et al., 2015). While these traits likely emerged as a mechanism to conserve energy in the absence of food, their specific contributions to starvation resistance are unknown. In addition, both sleep and feeding are developmentally plastic behaviors, and are modulated by both shared and independent neural mechanisms during the larval and adult stages (Itskov and Ribeiro, 2013; Koh et al., 2006; Melcher and Pankratz, 2005; Pool and Scott, 2014; Szuperak et al., 2018). *Drosophila* eat voraciously throughout development, and this is essential for organismal growth and the generation of energy stores that persist through adulthood (Merkey et al., 2011; Tennessen and Thummel, 2011). In addition, we have recently characterized larval sleep and found that this sleep is critical for development (Szuperak et al., 2018). Therefore, it is possible that selection for starvation resistance differentially influences adult behavior and physiological function, or that shared genetic architecture between development and adulthood results in an evolutionary constraint on developmental state-specific modification of behavior.

Here, we investigate sleep, feeding, and metabolic function throughout development in flies selected for starvation resistance. We find that development time is extended, starting at the 2^nd^ instar larval stage, and persists throughout development. In addition, whole-body metabolic rate is reduced during both development and adulthood and is accompanied by an increase in mass, suggesting that reduced energy expenditure allows these starvation resistant populations to increase their energy stores. Our findings also reveal that increased sleep and reduced feeding are specific to the adult stage, suggesting that selection for starvation resistance can target specific behaviors at different developmental time points.

## RESULTS

### Selection for starvation resistance in *Drosophila*

To assess developmental correlates of increased starvation stress, we utilized outbred populations that were artificially selected for starvation resistance. Briefly, flies were selected for starvation resistance by placing adult flies on agar and passaging starvation-resistant populations onto food when only ∼15% of flies remained alive. Three parallel starvation resistant groups were generated (S_A_, S_B_, and S_C_) as well as three controls that were continuously passaged on food (F_A_, F_B_, and F_C_). Experiments in this study utilized flies maintained on this selection protocol for 110-115 generations (Fig. 1A). In agreement with previous studies performed on flies selected for <60 generations (Hardy et al., 2018; Masek et al., 2014), this selection protocol robustly increased starvation resistance. All three S populations survived on average 9-13 days on agar compared to 2-3 days for the F populations (Fig. 1B,C), confirming that selection for starvation resistance results in approximately a four-fold increase in survival under starvation conditions.

**Fig. 1.**
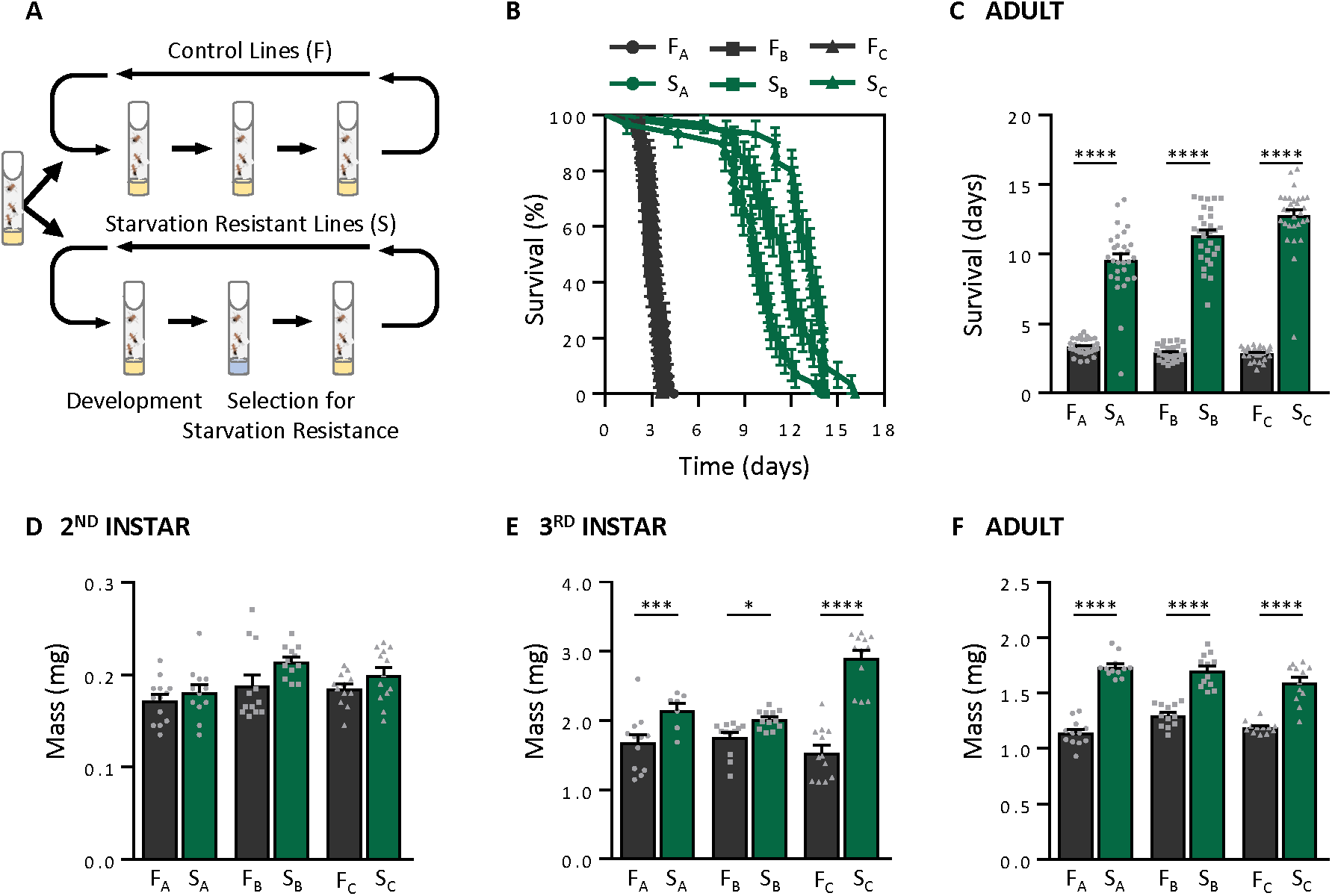
Selection for starvation resistance is correlated with larger body size. (A) Flies were selected for starvation resistance by maintaining adult flies on agar until ∼15% of flies remained alive. Three starvation resistant groups were generated as well as three fed control groups. (B,C) The S populations S_A_, S_B_, and S_C_) survive significantly longer on agar than the F populations (F_A_, F_B_, and F_C_) (Log-Rank test: χ^2^=210.7, d.f.=1, *P*<0.001; S and F populations pooled). (B) Survivorship curves showing the percentage of flies remaining alive as a function of the duration of starvation. (C) Mean survivorship of the S and F populations. Survivorship was measured once flies were transferred to agar. N = 27-32 per population. (D) Selection for starvation resistance increases mass in 2nd instar larvae (two-way ANOVA: F_1,66_ = 6.52, *P*<0.0130, N = 12 per population). However, post hoc analyses revealed no significant differences among replicate populations (A: *P*<0.8937; B: *P*<0.0681; C: *P*<0.3828). (E) Selection for starvation resistance increases mass in 3rd instar larvae (two-way ANOVA: F_1,66_ = 83.7, *P*<0.0001, N = 12 per population) and occurs in all three replicate populations (A: *P*<0.0003; B: *P*<0.0186; C: *P*<0.0001). In addition, we foundthat measurements of mass in 3rd instar larvae were population-specific (F_2,66_ = 8.645, *P*<0.0005) and that there was a significant interaction between mass and population (F_2,66_ = 16.88, *P*<0.0001). (F) Selection for starvation resistance increases mass in adults (two-way ANOVA: F_1,66_ = 266, *P*<0.0001, N = 12 per population), and occurs in all three replicate populations (A: *P*<0.0001; B: *P*<0.0001; C: *P*<0.0001). Similar to 3rd instar larvae, we found that measurements of mass in adults were population-specific (F_2,66_ = 4.954, *P*<0.0099) and there was a significant interaction between mass and population (F_2,66_ = 5.125,*P*<0.0001).

It has been previously shown that starvation-selection is associated with a larger body size in adult flies (Masek et al., 2014; Slocumb et al., 2015). To investigate whether this increase in body mass also occurs during development or is restricted to adults, we measured body mass during the 2^nd^ and 3^rd^ instar stages. Overall, we found that starvation-selected populations weighed significantly more than control populations at the 2^nd^ instar stage. However, we did not observe this effect when directly comparing each replicate F and S group individually (Fig. 1D). In the 3^rd^ instar stage, starvation-selected populations weighed significantly more than fed control populations, and each individual S group replicate weighed significantly more than their respective F control group (Fig. 1E). This increase in mass for all three starvation-selected replicate groups was maintained into adulthood (Fig. 1F). These findings suggest that starvation-selection is accompanied by an increase in mass that occurs during the 3^rd^ instar stage and persists through adulthood.

### Starvation-selection increases development time

It is possible that delayed development contributes to starvation resistance by allowing flies to accumulate energy stores during the larval stages. To determine whether the rate of development is altered by starvation-selection, we measured the time from egg laying to each developmental transition. Overall, development rate was delayed across all S groups, confirming that starvation-selection increases development time (Fig. 2A). A direct comparison of each developmental stage revealed no difference between F control groups and starvation-selected S groups in the transition from egg to first instar larvae, suggesting the selection protocol does not affect the earliest stages of development (data not shown). Development time was significantly delayed at all subsequent developmental stages (from 1^st^ instar to pupariation) when the three replicate starvation-selected populations and three control populations were each pooled. However, post hoc analyses on development time at each of these developmental stages revealed population-specific effects on the time spent within each stage. As such, a direct comparison of each replicate group revealed no differences in the duration of time spent as 1^st^ instar larvae (Fig. 2B). For the duration of time spent as 2^nd^ instar larvae, a similar comparison of each replicate group revealed that significant differences were only observed between the F_B_ and S_B_ populations (Fig. 2C). In contrast, all three S group replicate populations spent significantly longer during the 3^rd^ instar stage (Fig. 2D). During pupariation, significant differences in development time were again only observed between the F_B_ and S_B_ populations (Fig. 2E). Therefore, delayed development time is present across all starvation-selected populations, but is particularly robust in the S_B_ population. Overall, these findings raise the possibility that increased body size and starvation resistance are related to delayed development.

**Fig. 2.**
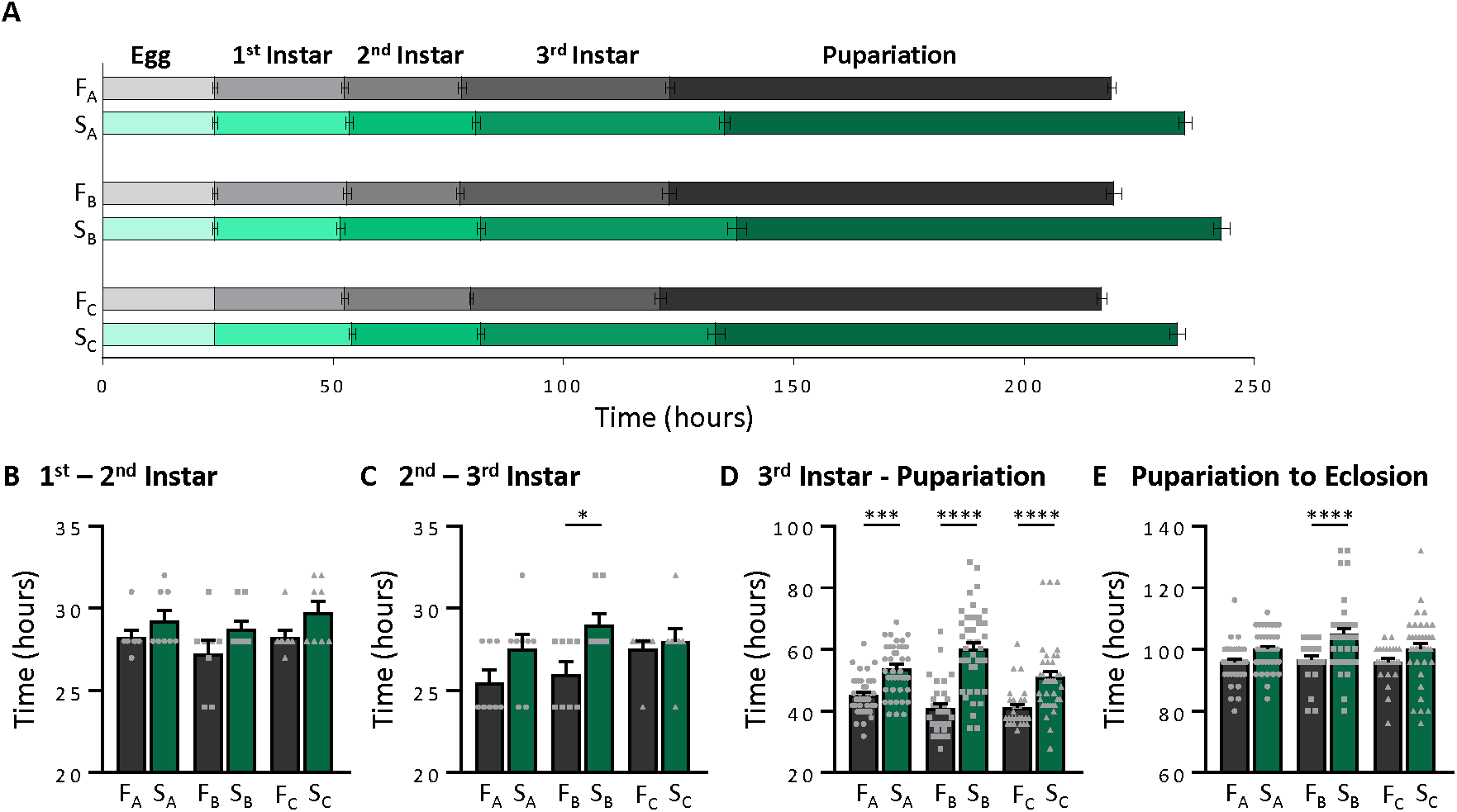
Selection for starvation resistance extends each stage of the *Drosophila* life cycle. (A) Average time spent in each stage of the life cycle from egg to eclosion. Increasingly darker bars indicate progression to later stages of larval and pupal development. (B) Average time it takes to molt from 1^st^ instar into 2^nd^ instar larvae. The S populations take longer to transition from 1^st^ instar to 2^nd^ instar larvae relative to the F populations (two-way ANOVA: F_1,42_ = 7.658, *P*<0.0084). However, post hoc analyses revealed no significant differences among replicate populations (A: *P*<0.5567; B: *P*<0.2199; C: *P*<0.2199). (C) Average time it takes to molt from 2^nd^ instar into 3^rd^ instar larvae. The S populations take longer to transition from 2nd instar to 3rd instar larvae relative to the F populations (two-way ANOVA: F_1,42_ = 9.517, *P*<0.0036). However, post hoc analyses revealed significant differences within the B population only (A: *P*<0.1661; B: *P*<0.0170; C: *P*<0.9492). (D) Average time it takes for 3^rd^ instar larvae to begin pupariation. The S populations take longer to transition from 3^rd^ instar into prepupae relative to the F populations(two-way ANOVA: F_1,217_ = 94.81, *P*<0.0001), and occurs in all three replicate populations (A: *P*<0.0001; B: *P*<0.0001; C: *P*<0.0001). (E) Average time from pupariation to eclosion. The S populations take longer to eclose from the pupal phase as adult flies relative to the F populations (two-way ANOVA: F_1,217_ = 27.07, *P*<0.0001), and occurs in all three replicate populations (A: *P*<0.0521; B: *P*<0.0001; C: *P*<0.0893). Egg-3^rd^ instar measurements: N = 8; pupation and eclosion measurements: N = 28-40.

### Starvation-selection decreases metabolic rate

In addition to delayed development, reduced metabolic rate provides a mechanism for conserving energy (Dulloo and Jacquet, 1998; Ma and Foster, 1986). Animals, including *Drosophila*, reduce metabolic rate under starvation conditions (Crabtree, 1990; McCue, 2010; Wang et al., 2006), suggesting that modulation of metabolic rate may promote starvation resistance. To determine the effect of starvation-selection on metabolic rate, we used indirect calorimetry to determine CO_2_ release, a proxy for metabolic rate, in both larvae and adults. Measurements of metabolic rate were then normalized to body mass in order to account for differences in body size between the F control groups and starvation-selected S groups. The system used to measure metabolic rate is highly sensitive, and has previously been used to detect CO_2_ release from single flies (Fig. 3A; Stahl et al., 2017a). In 2^nd^ instar larvae, no changes in metabolic rate were detected between F control groups and starvation-selected S groups (Fig. 3C). However, metabolic rate was reduced in 3^rd^ instar larvae when the three replicate starvation-selected populations and three control populations were each pooled. Post hoc analyses of each replicate population revealed a significant decrease in metabolic rate in the S_A_ and S_C_ populations compared to their respective F control populations (Fig. 3D). In adult flies, metabolic rate was significantly decreased in all three starvation-selected populations (Fig. 3E). Therefore, selection for starvation resistance results in reduced metabolic rate that commences during the 3^rd^ instar stage and persists into adulthood.

**Fig. 3.**
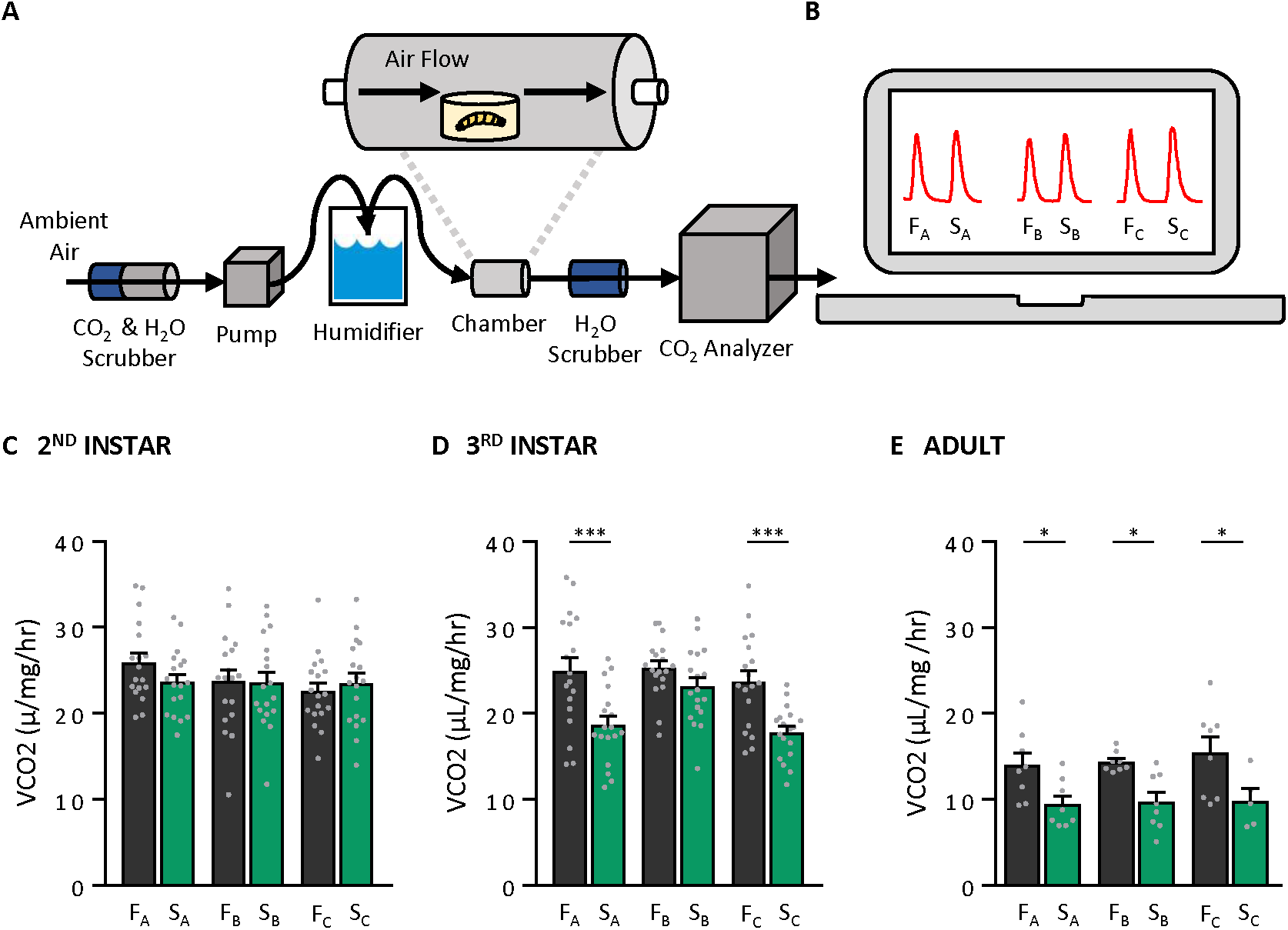
Selection for starvation resistance decreases metabolic rate and occurs in the later stages of larval development. (A) Metabolic rate was measured in 2^nd^ and 3^rd^ instar larvae and adults. Measurements were taken using a stop-flow respirometry system that measured the amount of CO_2_ produced over time. (B) Representative traces of each F and S population indicating the unadjusted amount of CO_2_ produced within each experimental chamber over time. (C) There is no change in metabolic rate in 2^nd^ instar larvae (two-way ANOVA: F_1,102_ = 0.3521, *P*<0.5543, N = 18 per population). (D) In 3rd instar larvae, selection for starvation resistance significantly decreases metabolic rate (two-way ANOVA: F_1,102_ = 0.27.89, *P*<0.0001, N = 18 per population). However, this effect is population specific (A: *P*<0.0004; B: *P*<0.4276; C: *P*<0.0008). (E) Metabolic rate is also significantly reduced in adults (two-way ANOVA: F_1,39_ = 21.71, *P*<0.0001, N = 4-6 per population), and persists in all replicate populations (A: *P*<0.0396; B: *P*<0.0318; C: *P*<0.0235).

**Fig. 4.**
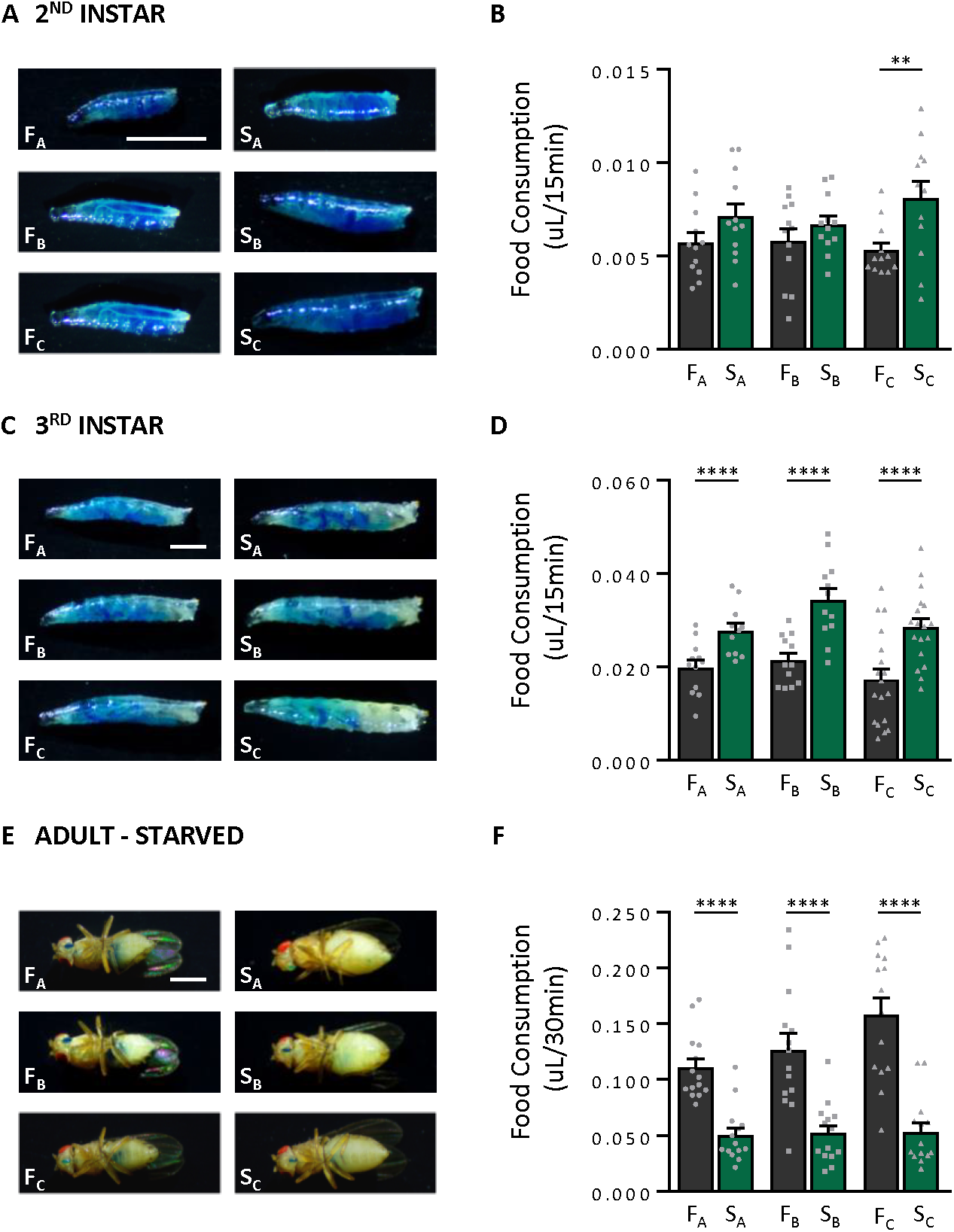
Selection for starvation resistance is correlated with an increase in food consumption beginning in the 3^rd^ instar stage. (A) Representative 2^nd^ instar larva from each population after 15 min of feeding on yeast paste supplemented with 2.5% blue dye. (B) Overall, starvation resistant populations consumed significantly more as 2nd instar larvae (two-way ANOVA: F_1,66_ = 10.68, *P*<0.0017, N = 12 per population). However, post hoc analyses revealed that only the S_C_ group increased food consumption relative to its control (A: *P*<0.3158; B: *P*<0.6912; C: *P*<0.0088). (C) Representative 3^rd^ instar larva from each population after 15 min of feeding on yeast paste supplemented with 2.5% blue dye. (D) Starvation resistant populations consumed significantly more as 3rd instar larvae (two-way ANOVA: F_1,71_ = 39.02, *P*<0.0001, N= 12-18 per population), and post hoc analyses revealed that this is the case for all three replicate groups (A: *P*<0.0062; B: *P*<0.0002; C: *P*<0.0001). (E) Representative starved adult female from each population after 30 min of feeding on food media supplemented with 2.5% blue dye. (F) Adults from starvation resistant populations consumed significantly less after 24hr of starvation than their respective controls (two-way ANOVA: F_1,78_ = 86.21, *P*<0.0001, N = 14 per population), and post hoc analyses revealed that this is the case for all three replicate groups (A: *P*<0.0004; B: *P*<0.0001; C: *P*<0.0001).

### Differential effects of starvation-selection on feeding and sleep

We have previously shown that food consumption is reduced in fasted starvation-selected adult flies (Masek et al., 2014). However, the effects of selection on larval feeding remain unknown. To quantify feeding in 2^nd^ and 3^rd^ instar larvae, we measured food intake by placing flies on yeast-paste laced with blue dye. The amount of food consumed over a 15-minute period was then measured based on spectrophotometric analysis of dye consumed during this time period. Food consumption was significantly increased among 2^nd^ and 3^rd^ instar larvae when the three replicate starvation-selected populations and three control populations were each pooled. However, during the 2^nd^ instar stage, post hoc analyses revealed that this effect was only significant in the F_C_ and S_C_ groups (Fig. 5A,B). During the 3rd instar stage, food consumption was increased in all three starvation-selected replicate populations (Fig. 5C,D),suggesting that starvation-selection promotes larval feeding. In contrast to larval feeding behavior, no differences were observed in food consumption across all three populations of starvation-selected adult flies in the fed state (Fig. S1). However, when animals were food deprived, so as to induce a robust feeding response, food consumption was significantly reduced across all three starvation-selected populations (Fig. 5E,F). These findings suggest that selection for starvation resistance has different effects on food consumption during the larval and adult stages.

**Fig. 5.**
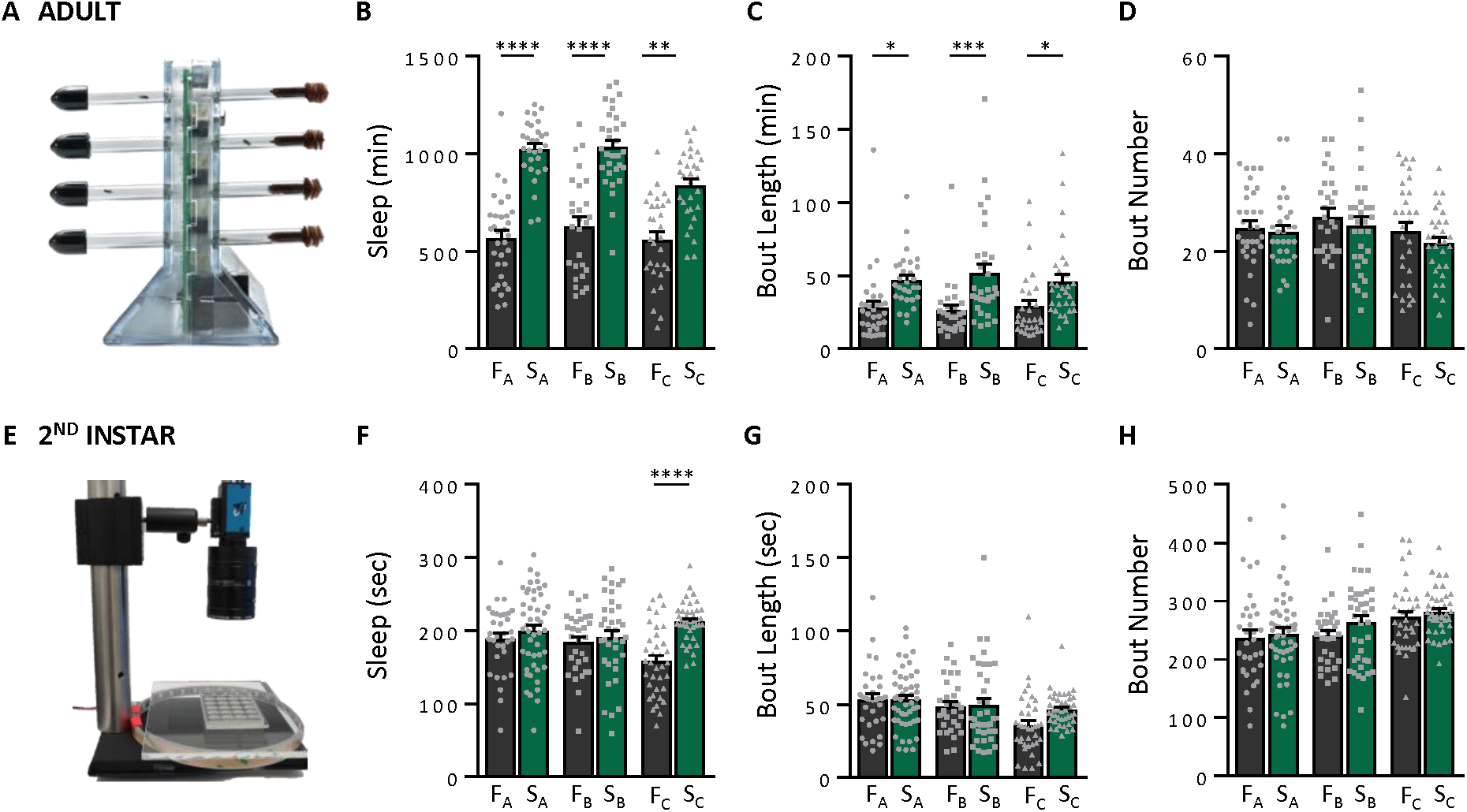
Selection for starvation resistance increases sleep duration and occurs in adults only. (A) Sleep traits in adults were measured using the *Drosophila* activity monitoring system. (B) Starvation resistant populations slept significantly more as adults (two-way ANOVA: F_1,173_ = 123.7, *P*<0.0001), and is consistent in all three groups (A: *P*<0.0001; B: *P*<0.0001; C: *P*<0.0021). The magnitude of the increase in sleep duration was population-specific, as there was a significant interaction between selection regime and line (F_2,173_ = 4.889, *P*<0.0087). (C) This increase in sleep duration is a result of an increase in the length of each sleep episode (two-way ANOVA: F_1,173_ = 29.87, *P*<0001), and is also consistent in all three groups (A: *P*<0.0110; B: *P*<0.0004; C: *P*<0.0288). (D) However, the number of sleep episodes does not differ (two-way ANOVA: F_1,173_ = 1.659, *P*<0.1994). Larvae: N = 31-48. Adults: N = 26-32. (E) Sleep traits in larvae were measured using custom-made LarvaLodges. (F) Starvation resistant populations slept significantly more as larvae (two-way ANOVA: F_1,220_ = 14.5, *P*<0.0002). However, post hoc analyses revealed that only the Sc population increased sleep relative to its control (A: *P*<0.6872; B: *P*<0.9047; C: *P*<0.0001). (G) The length of each sleep episode does not differ (two-way ANOVA: F_1,220_ = 2.351, *P*<0.1266), (H) nor does the number of sleep episodes (two-way ANOVA: F_1,1220_ = 2.304, *P*<0.1305).

It is possible that increased food consumption in starvation-selected larvae is a result of increased feeding drive or is secondary to their overall larger body size. To differentiate between these possibilities, we measured feeding rate by calculating the number of mouth hook contractions over a 30-second period. The number of mouth hook contractions did not differ between starvation-selected and control populations for 2^nd^ or 3^rd^ instar larvae (Fig. S2). These findings suggest that elevated food consumption in starvation resistant larvae results from increased food intake per mouth hook contraction and is likely related to their larger body size.

We previously reported that selection for starvation resistance increases sleep in adults (Masek et al., 2014). Here, we confirmed these results, finding that sleep duration was increased in all three starvation-selected populations, which is a consequence of increased bout length and not bout number (Fig. 5A-D). These results raise the possibility that energy conservation as a result of increased sleep may also occur during the larval stages. Recently, sleep has been characterized in 2^nd^ instar *Drosophila* larvae, allowing for the characterization of changes in sleep throughout development (Fig. 5E; Szuperak et al., 2018). Overall, we found that sleep increases among starvation-selected 2^nd^ instar larvae when the three replicate starvation-selected populations and three control populations were each pooled. However, post hoc analyses revealed that when sleep was assessed in each replicate population, an increase in sleep was only observed among the F_C_ and S_C_ populations, suggesting that these differences in sleep are present throughout development (Fig. 5F). No significant differences in bout length or bout number were detected between replicate populations of 2^nd^ instar larvae, though a trend towards increased bout length in the S_C_ population was detected, suggesting that sleep architecture is largely unaffected by selection for starvation resistance (Fig. 5G,H). Although we found no differences in sleep in 2^nd^ instar larvae, it is possible that additional sleep differences exist in 3^rd^ instar larvae; however, it remains unknown whether 3^rd^ instar larvae exhibit sleep states.

## DISCUSSION

Here, we report on the ontogenetically specified changes in behavior and metabolic rate induced by selection for starvation resistance. We found that starvation-selection extends larval development beginning in the 2^nd^ instar stage, with a concomitant decrease in metabolic rate and increase in food consumption beginning in the 3^rd^ instar stage. In adults, however, metabolic rate remains low, while food consumption remains unchanged and sleep is increased. These results suggest that starvation-selection has differential effects on behavioral and metabolic traits as development progresses from the larval stages into adulthood and is consistent with a strategy where starvation-selected larvae prioritize growth, while adults prioritize energy conservation.

Previous studies have found that selection for starvation resistance results in slower development in *Drosophila* (Chippindale et al., 1996; Hoffmann and Harshman, 1999; Masek et al., 2014; Reynolds, 2013), suggesting that extended larval development represents a mechanism for developing starvation resistance as adults. Here, we find that this reduced development rate begins as early as the 2^nd^ instar stage and persists until eclosion. During development, standard laboratory strains of *D. melanogaster* larvae increase their body size approximately 200-fold during the ∼4 days of larval development (Church and Robertson, 1966), raising the possibility that even subtle changes in development rate may significantly affect adult body size and energy stores. Progression through each larval transition is regulated by the steroid hormone 20-hydroxyecdysone, a master regulator of developmental timing (Riddiford, 1993; Liu et al., 2017; Yamanaka et al., 2013), and it is possible that starvation-selection acts to modify expression of this hormone, thereby delaying the onset of each larval transition. Additionally, several genetic factors have been identified that regulate nutrient dependent changes in developmental timing, including the target of rapamycin signaling pathway (Colombani et al., 2003; Layalle et al., 2008), and insulin-like peptides (Ikeya et al., 2002; Slaidina et al., 2009). Although significant advances have been made in elucidating the mechanisms underlying larval development and growth, our understanding of how environmental conditions, including starvation, can modulate these factors remain poorly understood. Our study reveals that environmental stressors can be potent selective forces that have a strong impact on the timing of larval growth and development.

It is proposed that animals develop resistance to starvation stress by reducing energy expenditure (Aggarwal, 2014; Hoffmann and Parsons, 1989; Marron et al., 2003; Rion and Kawecki, 2007). Here, we find that metabolic rate is reduced in both 3^rd^ instar larvae as well as in adults across all starvation-selected populations tested. While to our knowledge, the metabolic rate of *Drosophila* larvae has not previously been studied, earlier reports examining metabolic rate in adults from different populations of *D. melanogaster* selected for starvation stress found conflicting effects of selection on metabolic rate in flies (Baldal et al., 2006; Djawdan et al., 1997; Harshman and Schmid, 1998; Harshman et al., 1999; Marron et al., 2003). However, there is evidence that selection for starvation resistance results in altered use of metabolic enzymes in response to starvation (Harshman and Schmid, 1998), as well as an accumulation of energy stores (Masek et al., 2014; Schwasinger-Schmidt et al., 2012; Slocumb et al., 2015). In our study, we measured the metabolic rate of adult flies over a 24-hour period, thereby including any potential variation in the circadian effects of feeding, sleep, and metabolic rate. Alternatively, it is possible that the independent origins of the selected populations resulted in selection on metabolic rate-dependent and-independent pathways leading to enhanced starvation resistance. However, our finding that metabolic rate is reduced in multiple independent lines of starvation-selected populations suggests that these differences may be attributed to the initial outbred populations of flies used to derive starvation resistance.

Increased body size is a fitness-related trait that promotes tolerance to stress (Ewing, 1961). As such, environmental perturbation and food shortages may uniquely affect fitness depending on the developmental stage in which these selective pressures occur. The selection protocol used in this study selectively applied nutrient shortages during adulthood only, limiting selection pressures to traits that enhance adult starvation resistance. However, we found that starvation-selection increases body size as soon as the 2^nd^ instar stage and persists throughout the rest of development. Similarly, we identified larval-specific effects on food consumption and sleep. We found that food intake is increased in larvae but reduced in adults, and that sleep is reduced in adults but unchanged in larvae. These findings provide further evidence that selection for starvation resistance results in ontogenetically specified behavioral phenotypes. It has been previously shown that selective stresses imposed during development contribute to altered behavioral states as adults. In several *Drosophila* species, for example, thermal stress applied during larval development confers resistance to thermal stress in adulthood (Levins, 1969; Goto, 2000; Horu and Kimuro, 1998; Maynard Smith, 2005). Therefore, selection for starvation resistance during a defined developmental window can impact a variety of traits at multiple stages throughout development.

Although we found that changes in development rate, metabolic function, and sleep differ in starvation-selected populations, the genetic contribution of each trait to starvation resistance, especially at each stage of development, is unknown. We observed increased sleep and decreased starvation-induced feeding in starvation-selected adults, traits that are not present during larval development. Although these traits provide a potential mechanism for energy conservation in adult flies, in larvae, development rate slows, sleep does not differ, and food consumption actually increases. Furthermore, we found that metabolic rate is reduced in both larvae and adults of starvation-selected populations. Taken together, these findings support a model by which a slower development provides increased time to grow and accumulate energy stores as larvae, while reducing foraging-related behaviors in adulthood allows for animals to conserve energy as adults when food is not present during the selection process. This model suggests that distinct genetic architecture regulates sleep and feeding during the larval and adult stages. For instance, the mechanisms controlling larval sleep are partially distinct from that of adult sleep (Szuperak et al., 2018). Overall, these findings provide proof-of-principle for ontogeny-specific correlated behaviors during the applied starvation-selection process.

The identification of developmental, metabolic, and behavioral differences in *Drosophila* populations selected for resistance to starvation suggest that multiple mechanisms likely contribute to the etiology of starvation resistance. Towards this end, association mapping in *Drosophila* identified a wide range of genes associated with starvation resistance, including those that are known regulators of development, metabolism, and nutrient response (Harbison et al., 2004; Nelson et al., 2016). The complex genetic architecture underlying these traits, and their interrelationship, suggests the evolution of starvation resistance is likely to be highly pleiotropic, and it is possible that distinct mechanisms contribute to starvation resistance in the three replicate populations. Furthermore, a previous study examining the genetic divergence between these populations identified 1,796 polymorphisms that significantly differed between starvation-selected and control populations. These polymorphisms mapped to a set of 382 genes, including genes associated with a wide variety of metabolic and physiological processes (Hardy et al., 2018). While these studies provide an initial framework for identifying genetic factors regulating traits contributing to starvation resistance, a typical limitation of studying selected populations is a lack of accessible genetic tools that can be applied to validate the phenotypic contributions of single genes. The recent application of gene-editing approaches to outbred and non-*Drosophila melanogaster* populations raises the possibility of examining the contributions of these candidate genes in the future.

These findings reveal evidence of an ontogenetic shift associated with selection for starvation resistance in *Drosophila melanogaster*. This work highlights the contribution of several energy-saving traits that are modulated throughout development, including changes in metabolic rate, size, sleep, and food consumption to confer resistance to starvation. The development-specific differences in sleep and feeding of the starvation-selected populations set the stage for elucidating the genetic basis of starvation resistance ontogeny.

## METHODS

### *Drosophila* maintenance and Fly Stocks

Starvation-selected populations at generation 120 were obtained and then tested and maintained off selection for a maximum of 5 generations. All populations were grown and maintained on standard *Drosophila* media (https://bdsc.indiana.edu/information/recipes/bloomfood.html) and maintained in incubators (Percival Scientific, Perry, IA) at 25°C and 50% humidity on a 12:12 LD cycle. For larval experiments, adult flies were maintained in population cages with access to grape juice agar and yeast paste (Featherstone et al., 2009). Unless stated otherwise, eggs were collected from the cages within two hours of being laid and then transferred into petri dishes containing standard food at a constant density of 100 eggs per dish.

### Development time

Eggs were transferred into petri dishes containing standard food and green food coloring, which allowed for easier viewing of the larvae, at a density of 25 eggs per dish. Larvae were then scored every four hours for their transition through the 1^st^, 2^nd^, and 3^rd^ instar stages. The time at which at least 50% of the larvae within each petri dish have transitioned through each developmental stage was recorded. Time to pupariation and eclosion were measured independently from each larval instar stage. Eggs were collected within two hours of being laid and placed individually into glass test tubes, each containing 2mL of standard *Drosophila* media. Tubes were then scored every four hours.

### Feeding Behavior

Short-term food intake in adult flies was measured as previously described (Wong et al., 2009). Briefly, sets of five 3-4 day-old female flies were either transferred to vials containing a damp Kimwipe and starved, or maintained on standard food for 24 hrs. At ZT0, flies from both treatments were transferred to food vials containing 1% agar, 5% sucrose, and 2.5% blue dye (Federal Food, Drug, and Cosmetic Act, blue dye no. 1). After 30 minutes, flies were flash frozen and stored for subsequent analyses. For food consumption measurements in larvae, eggs were obtained as previously described. Eggs were transferred to petri dishes containing standard food at a larval density of 100 larvae per dish. Food consumption was measured at 60 and 96 hours after egg laying for 2^nd^ and 3^rd^ instar larvae, respectively. Short-term food intake in 2^nd^ and 3^rd^ instar larvae was performed as previously described (Kaun et al., 2007). Briefly, larvae were transferred to petri dishes containing a thin layer of 1% agar and yeast paste with 2.5% blue dye. After 15 minutes of feeding, larvae were collected and then washed in ddH2O three times. The larvae were then flash frozen in groups of 10 and 5 for 2^nd^ and 3^rd^ instar larvae, respectively. Each larval and adult sample was homogenized in 400 µL PBS and then centrifuged at 4°C at 13,000 rpm. The supernatant was then extracted and its absorbance at 655 nm was calculated using a 96-well plate absorbance spectrophotometer (Bio-Rad Laboratories, Inc.). Each sample was measured in triplicate. Baseline absorbance was determined by subtracting the absorbance obtained from flies/larvae not fed blue dye from each experimental sample. The amount of food consumed was then determined from a standard curve. To assess feeding rate in 2^nd^ and 3^rd^ instar larvae, the number of mouth hook contractions were counted (Shen, 2012). For each group, 2^nd^ or 3^rd^ instar larvae were placed onto a petri dish containing agar and yeast paste. After a 1-minute acclimation period, larvae were videotaped and the number of mouth hook contractions within a 30-second period were counted.

### Mass

For adults, 3-5 day old female flies were isolated and placed on fresh media for 24 hours, and then the mass of groups of 10 flies were determined. For 2^nd^ and 3^rd^ instar larvae, mass was measured at 60 and 96 hours after egg laying and was determined using groups of 10 and 20 larvae, respectively.

### Sleep and waking activity

In adults, Individual 3-5 day-old mated female flies were placed into tubes containing standard food and allowed to acclimate to experimental conditions for at least 24 hours. Sleep and activity were then measured over a 24hr period starting at ZT0 using the *Drosophila* Locomotor Activity Monitor System (DAMs) (Trikinetics, Waltham, MA) as previously described (Hendricks et al., 2000; Shaw et al., 2000). The DAM system measures activity by counting the number of infrared beam crossings for each individual fly. These activity data were then used to calculate bouts of immobility of 5 min or more using the *Drosophila* Sleep Counting Macro (Pfeiffenberger et al., 2010), from which sleep traits were then extracted. In larvae, sleep and activity was measured as described (Szuperak et al., 2018). Briefly, individual freshly molted 2^nd^ instar larvae were loaded into wells of custom-made PDMS microplates (LarvaLodges) containing 3% agar and 2% sucrose with a thin layer of yeast paste. LarvaLodges were loaded into incubators at 25°C and time-lapse images were captured every 6 seconds under dark-field illumination using infrared LEDs. Images were analyzed using custom-written MATLAB software, and activity/quiescence determined by pixel value changes between temporally adjacent images. Total sleep was summed over 6 hours beginning 2 hours after the molt to 2^nd^ instar. Sleep bout number and average sleep bout duration was calculated during this same period.

### Starvation resistance

The same flies used to measure sleep were also used to measure starvation resistance. Following 24hrs of testing on standard food, flies were transferred to tubes containing 1% agar (Fisher Scientific, Hampton, NH) and starvation resistance was assessed. The time of death was manually determined for each individual fly as the last bout of waking activity.

### Metabolic rate

Metabolic rate was measured though indirect calorimetry by measuring CO_2_ production (Stahl et al., 2017). Staged larvae were placed in groups of five (2^nd^ instar) or individually (3^rd^ instar) onto a small dish containing standard food media. Each dish was placed into a behavioral chamber where larvae were acclimated for 30 minutes, approximately the time required to purge the system of ambient air and residual CO_2_. Metabolic rate was then assessed by quantifying the amount of CO_2_ produced in 5 min intervals for 1 hour period. All experiments were conducted during ZT0-6 so as to minimize variation attributed to circadian differences in sleep, feeding, or metabolic rate. Metabolic rate in adults were assessed as described previously (Stahl et al., 2017). Briefly, adult flies were placed individually into behavioral chambers containing a food vial of 1% agar and 5% sucrose. Flies were acclimated to the chambers for 24hrs and then metabolic rate was assessed by quantifying the amount of CO_2_ produced in 5 min intervals during the subsequent 24hrs. Metabolic data for each group were normalized for body weight by dividing metabolic rate by mass, measured as described above. All experimental runs included larvae/flies from a randomized order of starvation-selected and control populations, as well as a food-only control, to account for any variation between runs.

### Statistical analysis

To assess differences in survivorship between starvation-selected and control populations, starvation resistance was analyzed using a log-rank test. Log-rank tests were also used to assess differences in development time, from 1^st^ instar to eclosion. A two-way ANOVA was performed on measurements of metabolic rate, mass, food consumption, mouth hook contractions, and sleep traits (factor 1: selection regime; factor 2: replicate population). If significant differences were observed, Sidak’s multiple comparisons test was performed to identify significant differences within each replicate population. All statistical analyses were performed using GraphPad Prism 7.0 (GraphPad Software, La Jolla, CA).

Where represented, all figures bars indicate mean values, error bars indicate SEM, and gray shapes indicate individual data points.

## ACKNOWLEDGEMENTS

We thank members of the Keene lab for technical assistance and helpful discussions.

## AUTHOR CONTRIBUTIONS

Conceptualization: E.B.B., M.E.S., M.S., A.K., and A.C.K.; Methodology: E.B.B., M.E.S., M.S., and A.K.; Formal analysis: E.B.B., M.E.S., and M.S.; Investigation: E.B.B., M.E.S., M.S., A.K., and A.C.K.; Resources: A.G.G., M.K., and A.C.K.; Visualization: E.B.B., M.E.S., and M.S.; Writing - original draft: E.B.B. and A.C.K.; Writing - review & editing: E.B.B., M.E.S., M.S., A.K., A.G.G., M.S.K., A.C.K.; Supervision: M.K. and A.C.K.; Funding acquisition: M.K. and A.C.K.

## SUPPLEMENTARY INFORMATION

Supplementary information available online at XXX.

## SUPPLEMENTARY FIGURES

Supplementary Fig. 1. There is no change in food consumption among fed adults between starvation-selected and control populations. (A) Representative fed adult female from each population after 30 min of feeding on food media supplemented with 2.5% blue dye. (B) Starvation resistant populations consumed the same as control populations (two-way ANOVA: F_1,66_ = 1.996, *P*<0.1625, N = 12 per population).

Supplementary Fig. 2. Increased food consumption in starvation resistant larvae is not a result of changes in feeding rate. There is no difference in the number of mouth hook contractions taken during a 30 sec period in either (A) 2^nd^ instar larvae (two-way ANOVA: F_1,66_ = 0.0003, *P*<0.9870, N = 12 per population) or (B) 3^rd^ instar larvae (two-way ANOVA: F_1,66_= 1.569, *P*<0.2148, N = 12 per population).

